# Epitopedia: identifying molecular mimicry between pathogens and known immune epitopes

**DOI:** 10.1101/2021.08.26.457577

**Authors:** Christian A Balbin, Janelle Nunez-Castilla, Vitalii Stebliankin, Prabin Baral, Masrur Sobhan, Trevor Cickovski, Ananda Mohan Mondal, Giri Narasimhan, Prem Chapagain, Kalai Mathee, Jessica Siltberg-Liberles

## Abstract

Upon infection, foreign antigenic proteins stimulate the host’s immune system to produce antibodies targeting the pathogen. These antibodies bind to regions on the antibody called epitopes. Structural similarity (molecular mimicry) of epitopes between an infecting pathogen and host proteins or other pathogenic proteins the host has previously encountered can impact the host immune response to the pathogen and may lead to cross-reactive antibodies. The ability to identify potential regions of molecular mimicry in a pathogen can illuminate immune effects which are especially important to pathogen treatment and vaccine design. Here we present Epitopedia, a software pipeline that facilitates the identification of regions that may exhibit potential three-dimensional molecular mimicry between an antigenic pathogen protein and known immune epitopes as catalogued by the immune epitope database (IEDB). Epitopedia is open-source software released under the MIT license and is freely available on GitHub, including a Docker container with all other software dependencies preinstalled. We performed an analysis describing how various secondary structure states, identity between pentapeptide pairs, and identity between the parent sequences of pentapeptide pairs affects RMSD. We found that pentapeptides pairs in a helical conformation had considerably lower RMSD values than those in Extended or Coil conformations. We also found that RMSD is significantly increased when pentapeptide pairs are from non-homologous sequences.

## 1. Introduction

Pathogens present antigenic molecules that can elicit a host immune response. For proteins, an epitope is the portion of the antigen that is recognized and bound by an antibody. Occasionally, pathogen epitopes may resemble host epitopes, a phenomenon termed molecular mimicry. In instances of molecular mimicry, infection with a pathogen can trigger the production of antibodies that mistakenly cross-react with an epitope in a host protein, resulting in autoimmune complications (Cusick et al., 2012). Alternatively, molecular mimicry between two pathogens can offer protective immunity for both after infection with either one (Agrawal, 2019).

To the best of our knowledge there are currently no computational programs or pipelines readily available for the prediction of molecular mimicry of known epitopes, although programs exist to map peptides (mimotopes) onto the antigenic protein structure to identify a native epitope (Chen et al., 2012; Huang et al., 2008; Mayrose et al., 2007; Negi & Braun, 2009), to identify molecular mimicry in remote homologs (Armijos-Jaramillo et al., 2021), and to identify molecular mimicry in antibody-binding interfaces (Stebliankin et al., 2022).

We present Epitopedia, a computational pipeline for the prediction of molecular mimicry. Epitopedia identifies sequence and structural similarity between an antigenic protein of interest and experimentally verified linear epitopes found in the Immune Epitope Database (IEDB) (Vita et al., 2019). Given the structural similarity between these epitopes and the pathogenic protein, it follows that binding of the same antibody may be possible.

## 2. Epitopedia Implementation

### 2.1 Internal Database Generation

Epitopedia utilizes data from the Immune Epitope Database (IEDB) (Vita et al., 2019), the Protein Data Bank (PDB) (Berman et al., 2000), and, optionally, the AlphaFold Protein Structure Database (Varadi et al., 2022) for the human proteome (Tunyasuvunakool et al., 2021). The data are organized into four internal tables (IEDB-FILT, mmcif-seqs, EPI-3D, and 3D-DSSP) stored in a SQLite3 database. IEDB-FILT is derived from a reduced IEDB that only includes the necessary data (epitope sequence, epitope identifier, antigen source sequence, range, accession, organism, etc.) for epitope mimicry search, including the full-length antigen source sequences from the IEDB. Based on the epitopes with positive assays from IEDB-FILT, a database for BLASTP (referred to as EPI-SEQ) of linear epitope sequences (mean length of 13 residues) and associated taxonomic origin of the epitopes is generated. Sequences from all PDB structures and human AlphaFold models were extracted and stored in mmcif-seqs. To find structural representatives for the antigen source sequences from IEDB, a sensitive (s=7.5) MMseqs2 (Steinegger & Söding, 2017) many-against-many search of antigen source sequences against mmcif-seqs is performed and the results are stored in EPI-3D. For a structural representative to be included in EPI-3D, the MMseqs2 pairwise alignment between the antigen source sequence and the structure sequence must have at least 90% identity or 20% query coverage. Lastly, DSSP (Kabsch & Sander, 1983) is used to determine secondary structure and compute the accessible surface area (ASA) for every residue in each chain in EPI-3D and the results are stored in 3D-DSSP.

### 2.2 Searching for 1-Dimensional Molecular Mimics

The Epitopedia pipeline is executed with one or more PDB IDs as input. The protein sequence (seqres) is extracted from the input structure and used in a BLASTP search against EPI-SEQ. The BLASTP parameters evalue and max_target_seq are both set to 2,000,000 to avoid discarding hits due to large evalues or reaching the match limit, respectively. The BLAST hits are filtered to only include hits with regions containing 5 or more consecutive, identical amino acids between the query (input protein based on the PDB ID input) and subject (epitope). If a hit meets this requirement in more than one region, the regions are split into subalignments so that one epitope may have >1 region.

Further, to be considered molecular mimics, the regions must have at least 3 consecutive accessible amino acids with a relative accessible surface area (RASA) > 20%. Based on ASA from 3D-DSSP and the maximum allowed solvent accessibility (MaxASA) values per amino acid as defined in Wilke (Tien et al., 2013), RASA is calculated according to equation

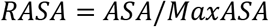

Regions meeting these qualifications are considered one dimensional mimics (1D-mimics). Regions that do not meet the aforementioned criteria to be considered a 1D-mimic are discarded.

### 2.3 Identifying 3-Dimensional Molecular Mimics

For 1D-mimics where the antigen source protein containing the epitope hit is represented in EPI-3D, the structural regions of the input structure corresponding to the 1D-mimic regions are evaluated to ensure that all residues are solved. To avoid missing potential mimics due to regions of missing electron density in an input structure, several structures can simultaneously be used as an input. Further, providing multiple PDB IDs for the same protein as input allows for a conformational ensemble approach to search for structural mimics. The structural fragments of 1D-mimics represented in EPI-3D and the corresponding hit fragment from the input structure are extracted. To compliment structural representation of human antigen source proteins in PDB, structural fragments can also be extracted from AlphaFold models for the human proteome (Tunyasuvunakool et al., 2021).

TM-align (Zhang & Skolnick, 2005) is used to evaluate the structural similarity based on the RMSD for each extracted peptide structure pair based on its BLAST hit pairwise alignment. To ensure that the structural superposition step is in agreement with the peptide pair sequence alignment, the pairwise alignment of the 100% identical 1D-mimic peptide pair is provided to TM-align. Pairs with an RMSD ≤ 1Å are considered three dimensional mimics (3D-mimics).

### 2.4 Handling Redundancy and Quantifying Results

Given the nature of epitopes and IEDB, it is common to have several overlapping epitopes where both the epitope mimic region and the antigen source sequence are identical. Internal accession numbers for all antigen source sequences in IEDB-FILT were assigned to ensure that any two or more identical sequences will have the same internal accession number to allow for filtering of redundancy at the output stage of the pipeline.

Epitopedia outputs results in CSV, JSON, and a simple web interface. The web interface is built using Flask, Bootstrap, and NGL Viewer (Rose et al., 2018) and provides an interactive visualization of the 3D-mimic region in both the input and epitope-containing antigen source protein. For each run, with N inputs, the distribution of RMSD values for the 3D-mimics is plotted as a histogram, with grey lines denoting the points of −1, 0, 1 standard deviations. The RMSD for each hit is denoted with a red line in the RMSD histogram. The Z-score for the hit is also computed, allowing for a comparative assessment of the hit quality against other hits for a particular run. An additional score termed EpiScore is calculated by dividing the mimic length by the RMSD (length of alignment/RMSD) to emphasize the significance of longer mimics. For example, given several mimics of varying length with the same RMSD, a longer mimic would have a higher EpiScore than a shorter mimic. Further, the EpiScore can reflect a more notable hit for a longer mimic with a higher RMSD than a shorter mimic with a lower RMSD. Thus, a higher EpiScore represents a more remarkable hit. As for RMSD, the EpiScore distribution for each run, shown as a histogram with the hit in red, is included in the web interface.

### 2.5 User customization

For each provided input structure, the following main steps allow for customization of the run. For the BLASTP search in *Step 1* (Figure 1), the user can specify a taxonomy filter for a focused search. With the taxonomy filter, epitopes from the specified taxonomic id will be excluded from the search.

**Figure 1.**
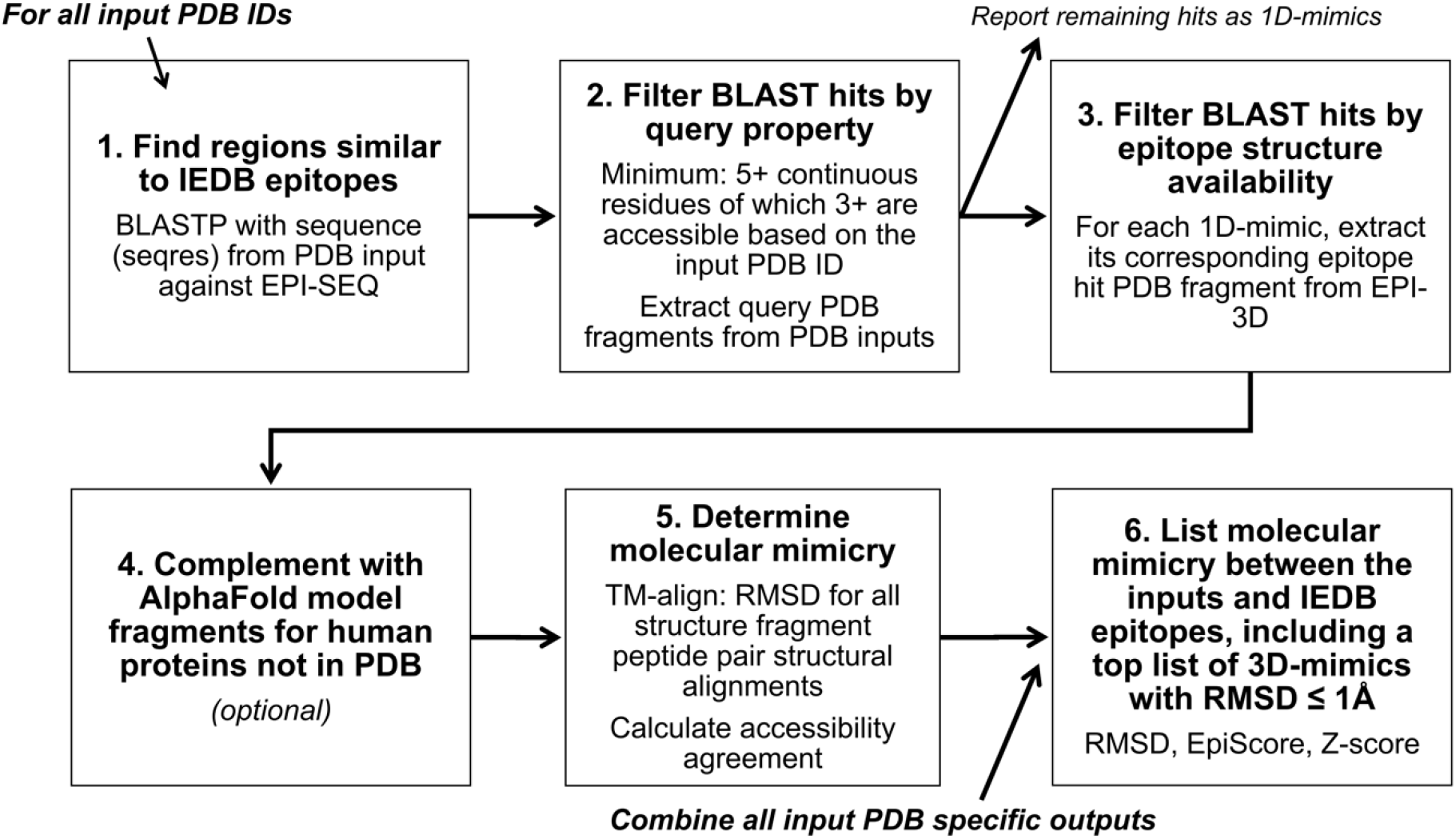
Overview of Epitopedia. Epitopedia is initiated with one or more PDB structures as input. In *Step 1*, a BLASTP search against linear epitope sequences in EPI-SEQ is performed with the corresponding sequence (seqres) from each PDB input as query. In *Step 2*, BLASTP hits that include sequence fragments from the query that do not contain at least 5 consecutive amino acids and where less than 3 amino acids are surface accessible based on the input structure are discarded. For the remaining hits, the PDB fragment is extracted from the input structure. These are considered 1D-mimics. In *Step 3*, structural fragments from the hits from EPI-SEQ that correspond to the 1D-mimics are extracted from PDB structural representatives of the source antigens. In *Step 4* (optional), for hits against epitopes in human source antigens that are not represented in PDB, structural fragments are extracted from AlphaFold models for regions with a certain confidence level (specified by the user). In *Step 5*, TM-align is used to calculate the RMSD of the structural alignment of the BLAST hit fragment or peptide pairs. In *Step 6*, RMSD results for all fragment pairs for all inputs for the run are combined. EpiScore (length of alignment/RMSD) and RMSD histograms are generated, and Z-scores are calculated based on the whole run. A top list of fragment pairs with RMSD ≤ 1Å is created. These fragment pairs are referred to as 3D-mimics.

For extracting potential epitope hits based on the input structure in *Step 2*, the minimum span length of an identical hit and the minimum accessibility of the hit in the input structure can be specified, with default values set to 5 and 3 residues, respectively. The user determines the cutoff for RASA, with the default set to 0.2. The sequence motifs from the epitope hits that meet span length and accessibility cutoffs are considered 1D-mimics, because although they are valid epitope hits based on the input structure, the structure of the epitope hit fragment is yet unknown. The structural fragments corresponding to the motif of each 1D-mimic are excised from the input structure.

In *Step 3*, for epitope hits corresponding to 1D-mimics from *Step 2*, the PDB structure of their antigen source protein is extracted from EPI-3D, if such a structure exists. Fragments matching the motifs of the 1D-mimics are excised for later comparison to the corresponding motif of each 1D-mimic from the input structure. Further, accessibility of the residues in the motifs is extracted from 3D-DSSP based on the whole protein structure.

Similarly, the user can choose to extract representative structures from an AlphaFold model of the human proteome (Tunyasuvunakool et al., 2021) based on EPI-3D in *Step 4*. The user can specify the confidence level of the AlphaFold models to consider using a motif (local) and a protein (global) confidence score. Both scores are based on pLDDT, which is the primary confidence score reported for AlphaFold models (Jumper et al., 2021). For the motif confidence score (m-pLDDT), no residue within the 1D-mimic motif can be below the cutoff. For the protein confidence score (p-pLDDT), the average of pLDDT for the entire model cannot be below the cutoff. The defaults are set to 0.9 and 0.7 for m-pLDDT and p-pLDDT, respectively. Structural fragments matching the motifs of the 1D-mimics are excised for later comparison to the corresponding motif of each 1D-mimic from the input structure. Further, accessibility of the residues in the motifs is extracted from 3D-DSSP based on the whole AlphaFold model.

In *Step 5*, structural comparisons of each motif fragment from the input structure to the corresponding fragments from *Step 3* or *Step 4* are performed using TM-align for the exact pairwise sequence alignment (Zhang & Skolnick, 2005). TM-score and RMSD are reported. However, because only short structural fragments are compared, the TM-score is not meaningful, while the RMSD of the structural alignment and agreement in RASA (based on the whole structural context) are meaningful. The user can set an RMSD cutoff for hits to be reported but the default is no RMSD cutoff.

In *Step 6*, all results for all input structures are compiled into a list. The EpiScore and Z-scores are computed. Hits with RMSD of at most 1Å are considered 3D-mimics. For the 3D-mimics, a web interface output is generated. The web interface includes the settings used to execute Epitopedia and basic information about the motif in the input structure, the epitope it mimics, and the antigen source protein in addition to RMSD, accessibility, EpiScore, Z-scores, and a link to a visualization of the results (Figure 2). For motifs with a 3D-mimic, the best hit is shown but the other hits are included under a dropdown menu. Structural visualization of 3D-mimics highlights the location of each mimic in the input structure and in the antigen source structural representative (Figure 3).

**Figure 2.**
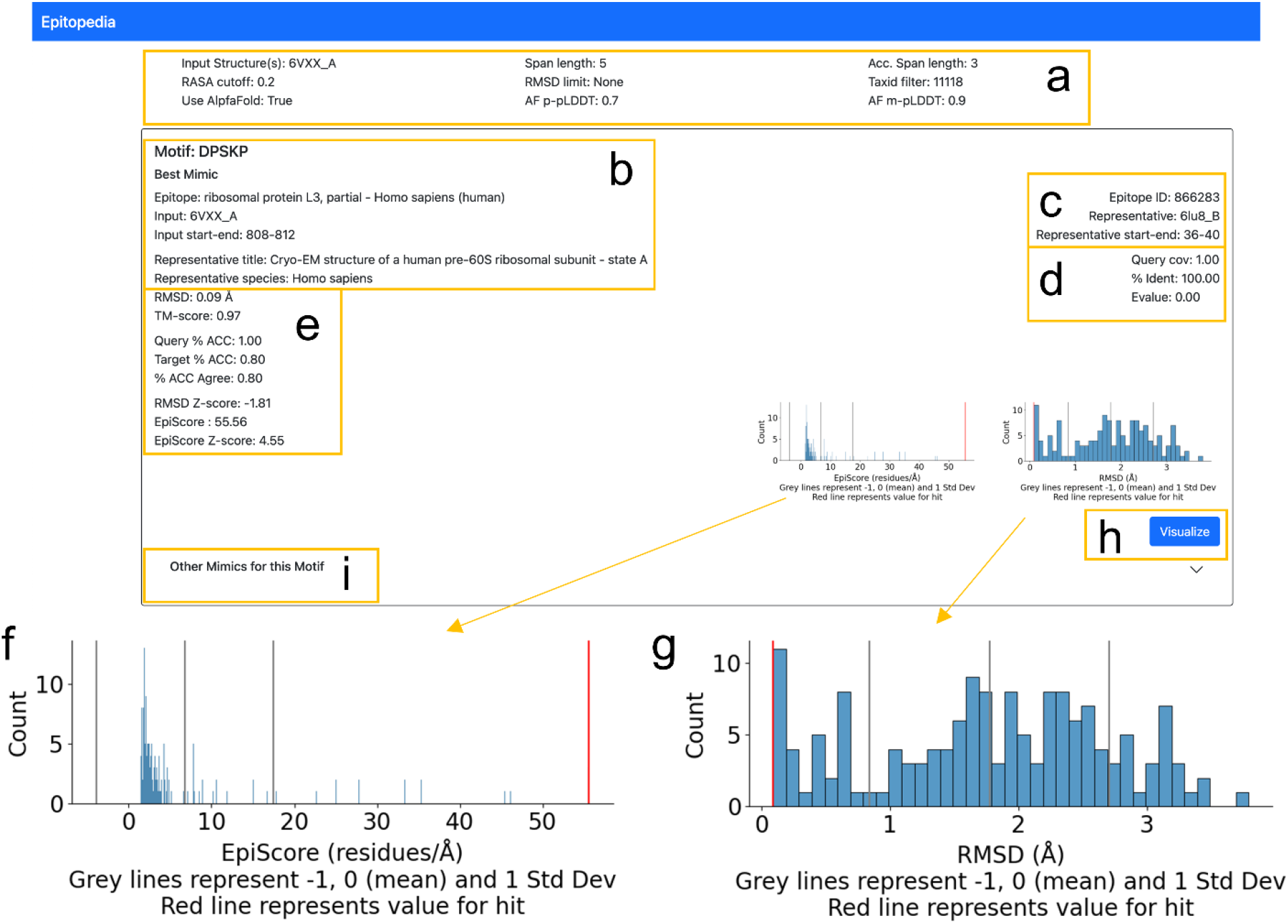
Overview of the Epitopedia web interface for 3D-mimics. For each run, (a) information about the run; (b) the mimic and protein in which the mimic was identified; (c) the epitope and its structural representative; (d) identification of the structural representative with MMseqs2; (e) structural comparison of the mimics including EpiScore, EpiScore Z-Score, and RMSD Z-Score; (f) EpiScore distribution for all structurally represented mimics (blue) during the given run including the EpiScore Z-score (grey), with the current mimic in red; (g) RMSD distribution for all structurally represented mimics (blue) during the given run including the RMSD Z-score (grey), with the location of the current mimic in red; (h) link to 3D visualization of the mimic; (i) and while the Best Mimic is shown from the start, additional mimics for the same motif from the same or different proteins but with higher RMSD are included in a dropdown menu.

**Figure 3.**
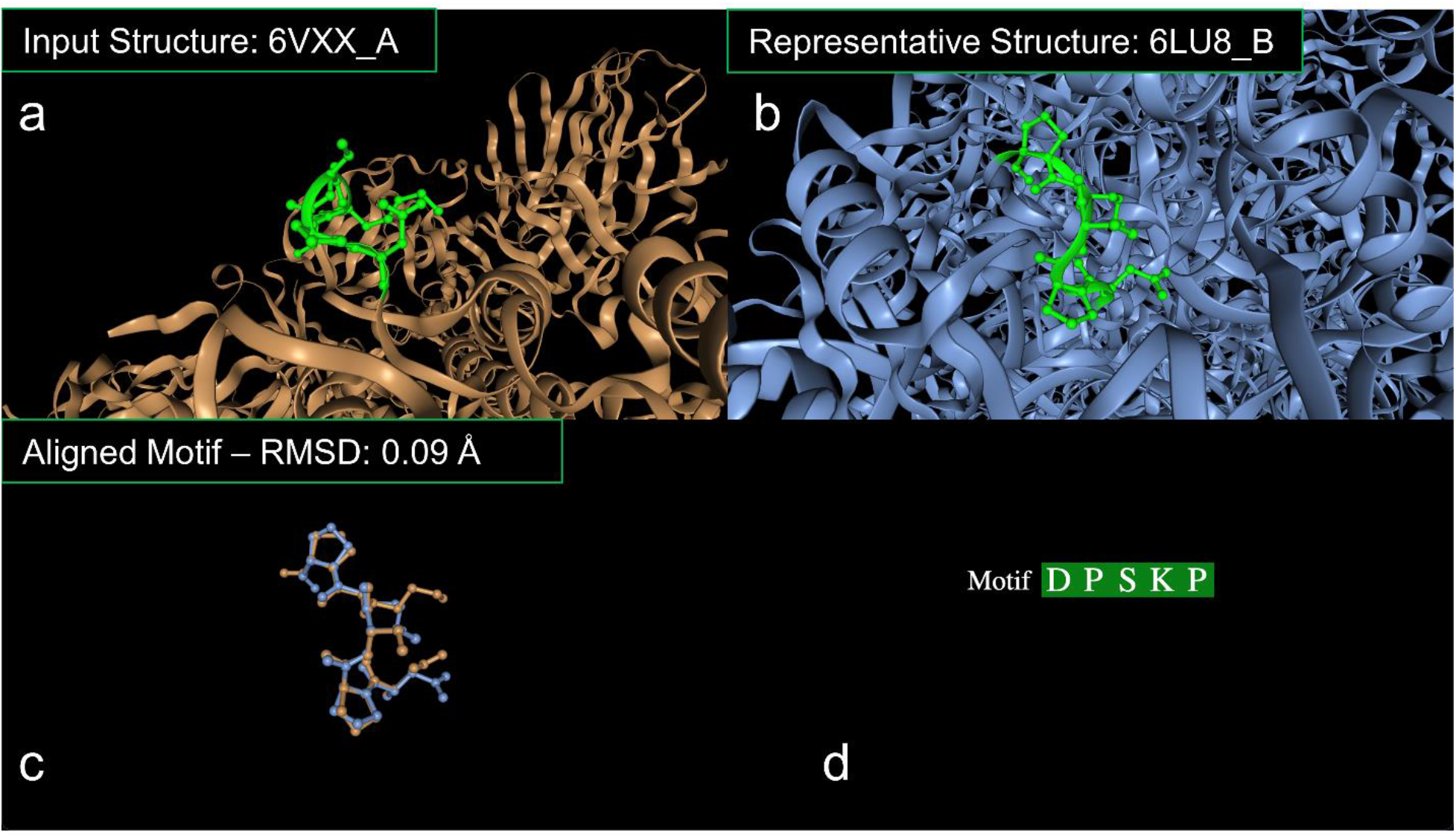
Visualization of the mimic pair in 3D. (a) The motif (green) shown in input protein (brown) (b) and in the structural representative protein (blue). (c) The TM-align structural superimposition for the motif in the input protein (brown) and the structural representative (blue). Panels a-c are interactive. (d) The mimic motif is interactive, hovering over a residue in the motif will highlight in panels a-c.

## 3. Epitopedia Demonstration

To demonstrate an Epitopedia run, we provide an example using an electron microscopy structure of the SARS-CoV-2 Spike protein (PDB ID: 6VXX, chain A (Walls et al., 2020)) as input. The taxid-filter flag with a taxid of 11118 was utilized to ensure neither the input protein nor other Coronavirus proteins were included as mimics (since these are homologous proteins and not mimics). The search for mimic representatives was performed against both PDB and the Human AlphaFold Protein Structure Database with default settings.

The run resulted in 755 1D-mimics, where 297 1D-mimics are structurally represented, of which 93 are only represented in the Human AlphaFold Protein Structure Database. After ensuring that only the best mimic per source sequence progresses, there were 153 mimics, with 66 of them mimicked with an AlphaFold structure. Finally, after filtering the results so that only 3D-mimics with an RMSD ≤ 1Å remain and removing redundant hits, there were 27 mimics, of which 11 are mimicked with an AlphaFold structure. Of the 16 3D-mimics from PDB, 13 are from human (such as integrin beta-1), and one each are from *Mycobacterium tuberculosis*, *Bacillus anthracis*, and Timothy grass (Table S1, Figures S1–S6). The remaining 11 3D-mimics are from the Human AlphaFold Protein Structure Database (Tunyasuvunakool et al., 2021; Varadi et al., 2022) and thus are all from human epitopes (Table S2). The mimic with the lowest RMSD (0.09 Å) is shown in Figures 2 and 3.

We also applied Epitopedia to a different SARS-CoV-2 Spike structure (PDB ID: 6XR8) and identified additional molecular mimicry with potential implications for COVID-19 (Nunez-Castilla et al., 2021).

## 4. Pentapeptide Structural Space Analysis

To provide guidance on how to interpret structural mimicry based on RMSD for the 3D-mimic pentapeptide fragment pairs identified by Epitopedia, we performed an investigation of RMSD for random pentapeptide pairs for the three main secondary structure states helix, extended, and coil from any sequence pair regardless of sequence similarity and for sequence pairs with low sequence similarity representing non-homologous proteins.

### 4.1 Methods

To understand how secondary structure state and sequence identity affect the distribution of RMSD values for pentapeptide pairs, an analysis of RMSD distributions of pentapeptide pairs across various secondary structure states and pentapeptide sequence identity levels was performed.

All possible pentapeptides based on PDB structures were generated and annotated with a DSSP secondary structure state reduction based on 3D-DSSP. The DSSP state reduction was performed such that if all residues in a pentapeptide were classified as turn (T), bend (S) or none (-), the pentapeptide was labeled coil, if all residues were strand (E) or beta-bridge (B) the pentapeptide was labeled extended, and if all residues were alpha helix (H), 3-10 helix (G), or pi-helix (I) the pentapeptide was labeled helix. Any pentapeptides that did not fit into one of these 3 categories were discarded.

Around 1,000 pentapeptide pairs (Table S3) were generated per secondary structure per identity level (0%, 20%, 40%, 60%, 80%, and 100%) from the labeled pentapeptide database described above. The number of pentapeptide pairs per category is not exactly the same across all categories because matches of a pentapeptide against itself (same PDB ID) are discarded. The pentapeptide regions were extracted from the parent structures using GEMMI (*GitHub - Project-Gemmi/Gemmi: Macromolecular Crystallography Library and Utilities*, n.d.) and superposed using TM-align (Zhang & Skolnick, 2005), with a fixed alignment as described for the Epitopedia implementation above.

To reduce the influence that parent sequence homology may have on the above analysis, we performed a similar analysis starting with 2,000 pentapeptides per secondary structure per identity level. Here, an added filtering step was performed to ensure that the parent sequences of the pentapeptide pairs were no more than 30% identical according to a local pairwise Smith-Waterman alignment of the parent sequences generated with EMBOSS Water (Madeira et al., 2019). Pentapeptide matches where the identity filter could not be enforced were discarded, thus, the number of pentapeptide pairs per category is not exactly the same across categories. For instance, if a query pentapeptide had been paired with over 100 other pentapeptides to generate a pentapeptide pair, yet a pentapeptide pair with a parent sequence identity of less than 30% was not found, the query pentapeptide was discarded. This scenario disproportionately affected pentapeptide pairs with higher pentapeptide identity, as there is a lower chance of parent sequences having less than 30% identity as the pentapeptide pair identity increases. In total, all pentapeptide identity and secondary structure combinations have greater than 900 pentapeptide pairs (Table S3).

Statistical comparisons were performed with Mann Whitney U using *SciPy* (Virtanen et al., 2020). Alpha values were corrected for multiple comparisons using simple Bonferroni correction. For a confidence level of 99%:

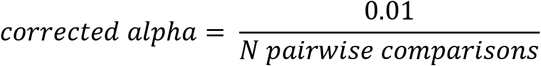

### 4.2 Results

An analysis was performed to better understand how the RMSD distribution for pentapeptides pairs varies with differing pentapeptide pair sequence identity, parent sequence identity and secondary structure state. For the first analysis that did not consider the percent identity of the parent sequences for a pentapeptide pair, a decrease in the median RMSD is observed at the 100% identity levels (Figure 5, Table 1). For helix pentapeptide pairs, the median RMSD for the 0% to 80% pentapeptide identity levels is 0.20-0.22Å, while at the 100% identity level the median is 0.13Å, which is a significant decrease when compared to all other identity levels for the helical state (Table 2). For extended pentapeptide pairs, the median RMSD for the 0% to 80% pentapeptide identity levels is 0.69-0.84Å, while at the 100% identity level the median is 0.14Å. This is a significant decrease when compared to all other identity levels for the extended state (Table 2). Lastly, for coil pentapeptide pairs, the median RMSD for the 0% to 80% pentapeptide identity levels is 1.79-1.95Å, while at the 100% identity level the median is 0.31Å. This large decrease of ~1.5Å is significant when compared to all other identity levels for coil (Table 2).

**Figure 4.**
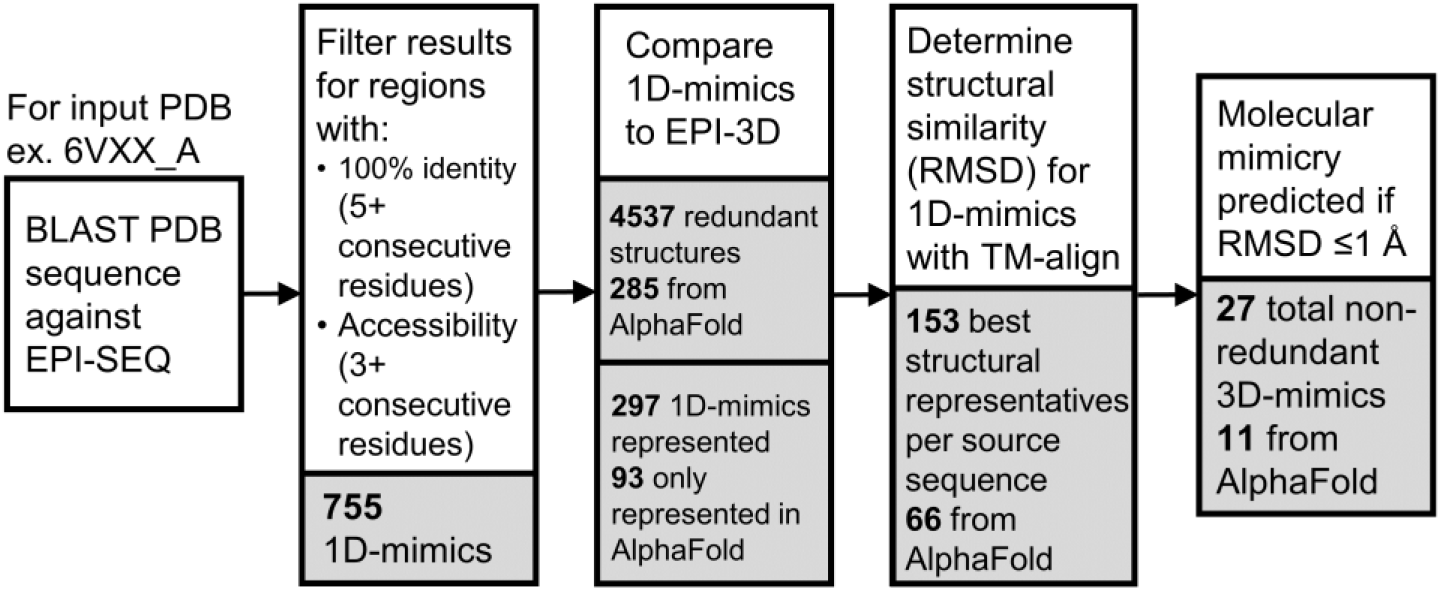
Epitopedia output overview using PDB 6VXX, chain A as input. For detailed output see example_output folder on the GitHub repository.

**Figure 5.**
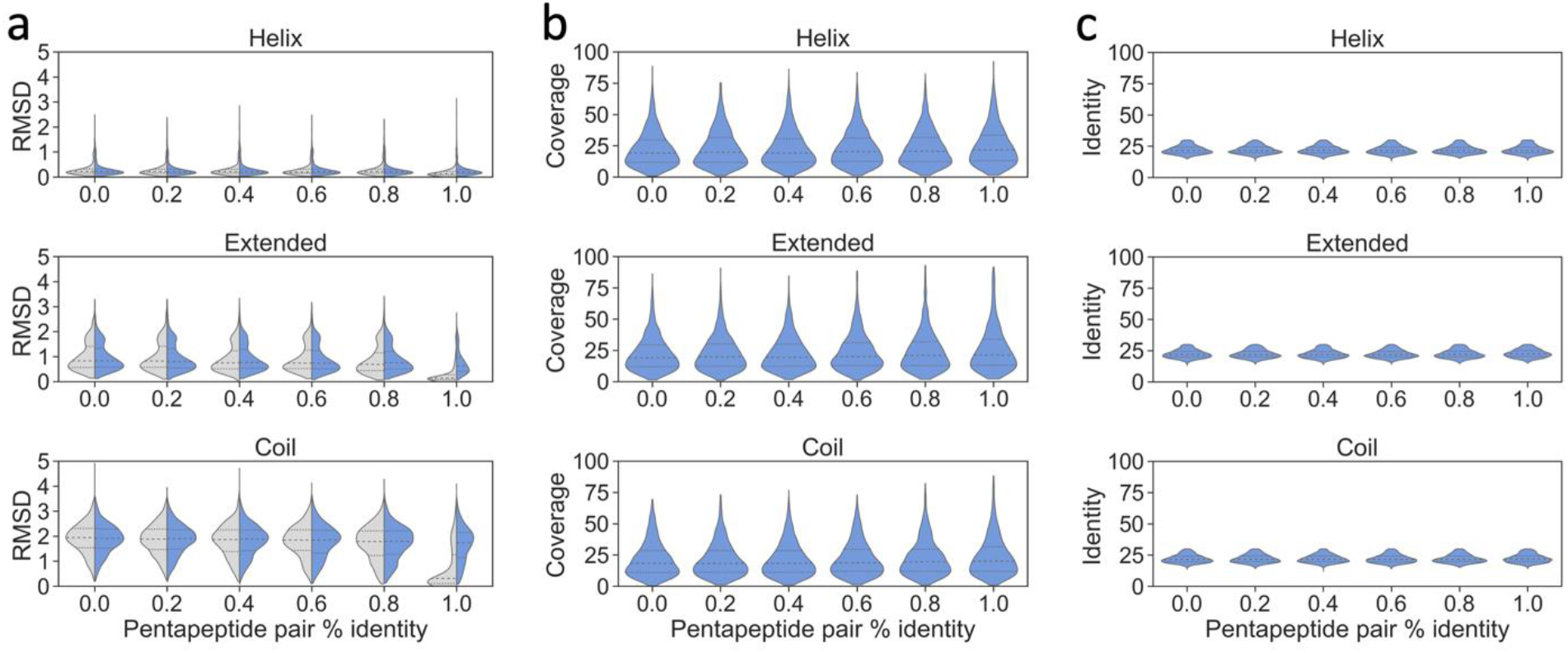
(a) Violin plots of the resulting RMSD distribution from pentapeptide structure analysis. The distributions for the analysis without the 30% parent sequence identity filter are shown in grey while the corresponding distributions for the pentapeptides from the 30% parent sequence identity set are shown in blue. (b) Violin plots showing the distribution of query coverage between the parent sequences for pentapeptide pairs at various identity levels and secondary structure categories. (c) Violin plots showing the distribution of pairwise identity between the parent sequences for pentapeptide pairs at various identity levels and secondary structure categories

**Table 1.** Median RMSD values resulting from RMSD distribution for structural space analysis of pentapeptide pairs of various identity levels and secondary structural categories shown in Figure 5.

| Pentapeptide<br>% Identity | Median RMSD (Å) |  |  |  |  |  |
| --- | --- | --- | --- | --- | --- | --- |
|  | No Parent Identity Filter |  |  | 30% Parent Identity Filter |  |  |
|  | Helix | Extended | Coil | Helix | Extended | Coil |
| 0 | 0.22 | 0.84 | 1.95 | 0.22 | 0.83 | 1.91 |
| 20 | 0.21 | 0.82 | 1.88 | 0.21 | 0.80 | 1.90 |
| 40 | 0.21 | 0.75 | 1.87 | 0.21 | 0.77 | 1.88 |
| 60 | 0.20 | 0.74 | 1.85 | 0.21 | 0.74 | 1.85 |
| 80 | 0.21 | 0.69 | 1.79 | 0.21 | 0.76 | 1.79 |
| 100 | 0.13 | 0.14 | 0.31 | 0.20 | 0.63 | 1.74 |

**Table 2.** Comparisons across pentapeptide identity levels for the same structural class for the no filter and 30% filter datasets, respectively. Significantly different comparisons are shown in blue.^1^

| Pentapeptide % Identity |  | No Parent Identity Filter |  |  | 30% Parent Identity Filter |  |  |
| --- | --- | --- | --- | --- | --- | --- | --- |
|  |  | Helix | Extended | Coil | Helix | Extended | Coil |
| 0 | 20 | 0.197694 | 0.578304 | 0.070875 | 0.084507 | 0.107303 | 0.464566 |
| 0 | 40 | 0.377122 | 0.000931 | 0.001582 | 0.212098 | 0.002897 | 0.013426 |
| 0 | 60 | 0.076141 | 0.000368 | 0.000528 | 0.15162 | 1.64E-06* | 7.41E-06* |
| 0 | 80 | 0.313838 | 3.52E-12* | 5.1E-09* | 0.0284 | 9.37E-08* | 1.97E-10* |
| 0 | 100 | 8.4E-59* | 1.3E-212* | 5.5E-156* | 5.63E-06* | 2.77E-31* | 3.33E-17* |
| 20 | 40 | 0.688416 | 0.006825 | 0.145049 | 0.636335 | 0.170195 | 0.083497 |
| 20 | 60 | 0.634811 | 0.00257 | 0.092602 | 0.779538 | 0.001278 | 0.000172 |
| 20 | 80 | 0.791704 | 1.43E-10* | 2.84E-05* | 0.639867 | 0.000168 | 1.84E-08* |
| 20 | 100 | 3.63E-52* | 2.8E-210* | 2.3E-149* | 0.002963 | 4.31E-25* | 6.02E-15* |
| 40 | 60 | 0.362381 | 0.785088 | 0.797398 | 0.838454 | 0.06416 | 0.038889 |
| 40 | 80 | 0.891271 | 7.14E-05* | 0.006063 | 0.35334 | 0.016001 | 8.83E-05* |
| 40 | 100 | 1.27E-54* | 4.5E-203* | 1.9E-141* | 0.00072 | 2.59E-20* | 2.54E-10* |
| 60 | 80 | 0.457191 | 0.000238 | 0.011388 | 0.457478 | 0.571466 | 0.067424 |
| 60 | 100 | 1.06E-50* | 7.3E-200* | 2.5E-140* | 0.001357 | 1.51E-14* | 6.59E-06* |
| 80 | 100 | 3.08E-53* | 2.2E-174* | 1.5E-124* | 0.01095 | 5.00E-13* | 0.003258 |

<sup>1</sup> Based on simplified Bonferroni correction at 99% confidence level, corrected alpha = 0.000111.

For the follow-up analysis we enforced a maximum 30% parent sequence identity filter to better resemble molecular mimics from unrelated protein pairs. To ensure that the pentapeptide pairs were from proteins that are not closely related, we performed local alignments and extracted the percent sequence identity and query cover for each parent sequence pair. By design, no parent sequence pair has a pairwise sequence identity above 30%, with a median around 20% (Figure 5). The query cover for the parent sequence pair alignments is low, with a median of around 20% (Figure 5). For these pairwise sequence alignments with 20% sequence identity and query cover, we can assume that these are primarily non-homologous parent sequence pairs although some remote homologs may be included in this dataset.

For the pentapeptide pairs from these non-homologous sequence pairs, the sharp decrease in the median RMSD at the 100% pentapeptide identity level has faded for extended and coil conformations (Figure 5). For helix pentapeptide pairs, the median RMSD at the 0% to 100% pentapeptide identity level is 0.20-0.22Å. Only the 100% vs 0% identity level comparison yields significant difference for the helix pentapeptide pairs (Table 2). For extended pentapeptide pairs, the median RMSD at the 0% to 100% pentapeptide identity level is 0.63-0.83Å. For coil pentapeptide pairs, the median RMSD at the 0% to 100% pentapeptide identity level is 1.74-1.91Å. For extended and coil pentapeptide pairs, the 100% identity level is significantly different when compared against every other identity level except for one comparison, 80% vs 100% in the coil state (Table 2).

When comparing the same identity level for the pentapeptide pairs across the set with no parent sequence identity filter and the set with the 30% sequence identity filter, we found that the pairwise parent sequence identity has an impact on the RMSD for identical pentapeptide pairs in the helical state, but not for the less identical peptide pairs (Table 3). This pattern is shared for the coil state, but for the extended state, the pairwise parent sequence identity seems to impact RMSD for identical and 80% identical pentapeptides (Table 3).

**Table 3.** Comparisons between pentapeptide identity levels for the same structural class for the 30% filter vs the no filter dataset. Significantly different comparisons are shown in blue.^1^

| Pentapeptide % Identity |  | 30% vs no Parent Identity Filter |  |  |
| --- | --- | --- | --- | --- |
|  |  | Helix | Extended | Coil |
| 0 | 0 | 0.438284984 | 0.56859737 | 0.1896511 |
| 20 | 20 | 0.381648164 | 0.255207316 | 0.7826223 |
| 40 | 40 | 0.445432037 | 0.339928559 | 0.5969043 |
| 60 | 60 | 0.091388265 | 0.844626827 | 0.4058797 |
| 80 | 80 | 0.856800634 | 0.000269319* | 0.5128634 |
| 100 | 100 | 3.41E-55* | 4.83E-171* | 1.93E-131* |

<sup>1</sup>Based on simplified Bonferroni correction at 99% confidence level, corrected alpha = 0.000555556.

Altogether, this analysis shows that for pentapeptides, the secondary structure state is important to consider when identifying molecular mimics using RMSD for random proteins. We used TM-align to calculate RMSD and this method, like many others, calculates RMSD based on the spatial coordinates for C-alpha in each amino acid residue. Our observation that pentapeptide pairs in a helical state have lower RMSD is not surprising given the regular geometry of the α-helix. For identical pentapeptide pairs in extended and coil conformations, the median RMSD for the non-homologous parent sequences are 0.63Å and 1.74Å, respectively, compared to 0.20Å for helix (Table 1).

### 4.3 Guidelines

Our interpretation, as far as molecular mimicry goes, is that mimics with identical sequences in α-helices are likely to appear very similar if they are oriented the same way in their parent proteins. As such, they are likely to be able to participate in similar interactions with, for example, an antibody. Mimics with identical sequences with low RMSDs, approaching the median RMSD of the unfiltered set (Table 1), are likely to present a similar interaction interface, if oriented similarly. A pentapeptide in a helix, given its winding structure, is relatively short while a pentapeptide in the extended or coil conformation may present a larger accessible area.

Pathogen proteins that mimic known epitopes in antigenic proteins may stimulate the production of cross-reactive antibodies that can interact with the pathogen protein as well as the human antigen. Pathogen proteins that mimic known epitopes in other pathogens may trigger an immune memory that could lead to protective immunity or complex immune effects such as antibody dependent enhancement.

## 5. Conclusion

Here, we have developed Epitopedia, a pipeline for the discovery of potential molecular mimics of immune epitopes found in IEDB. Epitopedia can facilitate our understanding of how pathogens may interfere with the known epitopes from the human proteome and also known epitopes from other species. Epitopes shared between pathogens can impact immune responses for secondary infections and identification of mimics of epitopes can provide insights to the mechanism behind the widely differing clinical manifestations and complications of infection with certain pathogens, such as SARS-CoV-2. Identification of molecular mimicry between known epitopes from the human proteome and a human pathogen protein can provide clues to the autoimmune potential of an infection caused by the pathogen. Further, by pinpointing regions in the pathogen’s proteome that may cause an autoimmune response if a cross-reactive antibody is created against it, these regions can be avoided in future vaccine design. Lastly, by highlighting which human proteins may be at risk for autoimmune targeting in response to a pathogen infection, therapeutics to counteract autoimmune effects can be used (or developed). Epitopedia provides a starting point for generating a better understanding of the autoimmune potential of pathogens and can benefit large-scale data mining and experimental in-vitro and in-vivo design to solve autoimmune conundrums in infectious disease.

## Acknowledgments

We thank Teresa Liberatore for discussions.

## Author Contributions

J.S.-L., G.N., P.C., K.M., T.C., A.M.M. conceived the overall approach. J.S.-L., J.N.-C. conceptualized the Epitopedia pipeline. J.N.-C., C.A.B., J.S.-L. designed and developed the method and approach. J.N.-C. created a pilot of the core pipeline. C.A.B. wrote the code to implement the pipeline. J.N.-C., C.A.B., J.S.-L. analyzed data and performed visualization. J.S.-L supervised the project. All authors discussed the pipeline and provided feedback. J.N.-C., C.A.B., J.S.-L. wrote the manuscript. All authors read and commented the manuscript.

## Data and Software Availability

Epitopedia is primarily written in Python and relies on established software and databases. Epitopedia is available at https://github.com/cbalbin-bio/Epitopedia under the opensource MIT license and also as a docker container at https://hub.docker.com/r/cbalbin/epitopedia.

## Supplementary Materials

Supplementary tables S1–S3 and figures S1–S6

## SUPPLEMENTARY MATERIALS

**Table S1.**
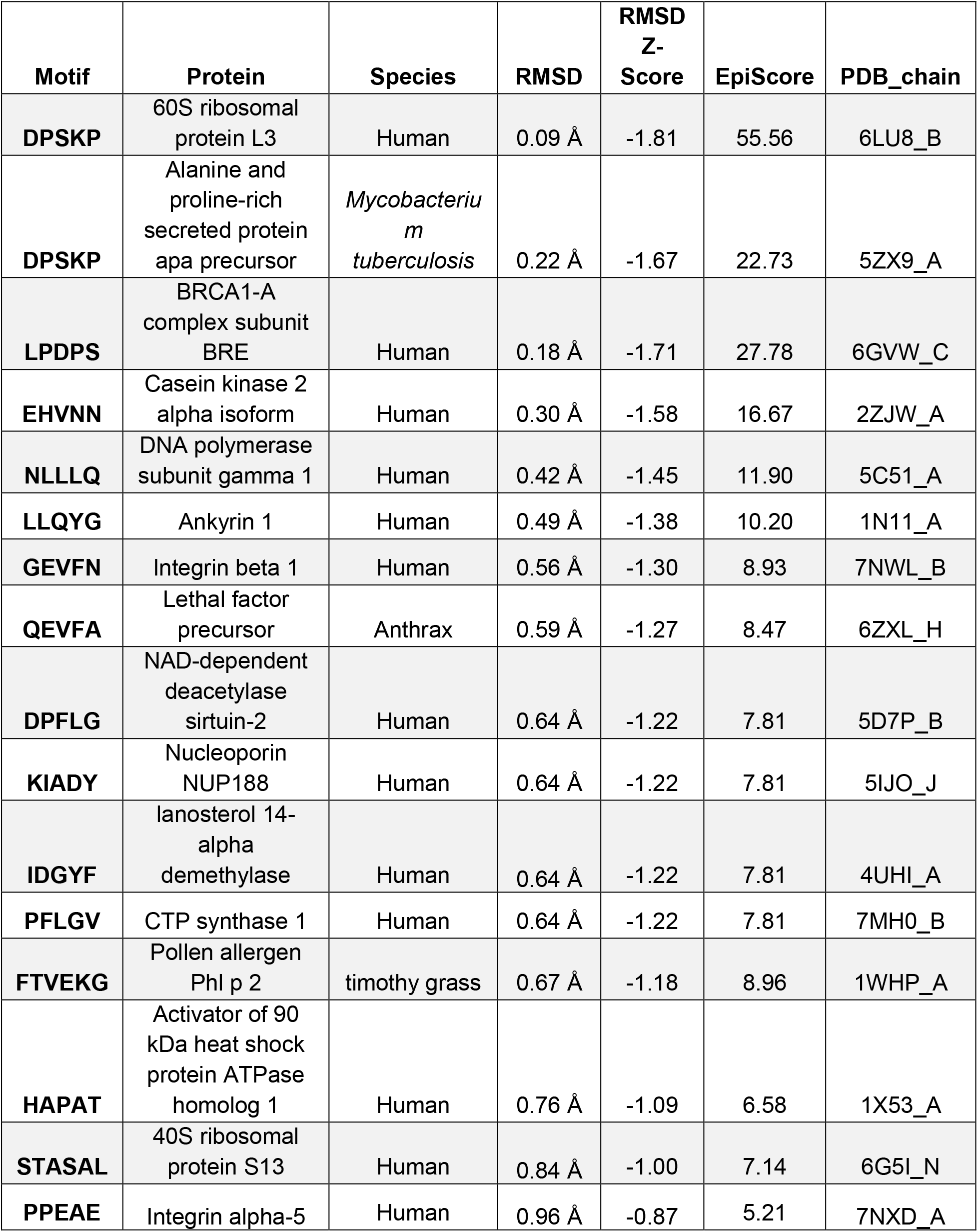
3D-mimics found for SARS-CoV-2 Spike (PDB id 6VXX_A)

**Table S2.** Human AF-3D-mimics for SARS-CoV-2 Spike

| **Motif** | **Protein** | **RMSD** | **RMSD Z-Score** | **EpiScore** | **AlphaFold2 ID** |
| --- | --- | --- | --- | --- | --- |
| **TGIAV** | Phosphofructokinase | 0.09 Å | -1.81 | 55.56 | AF-P17858-F1-model_v1_A |
| **TGIAV** | Low affinity immunoglobulin gamma Fc region receptor II-b | 0.11 Å | -1.78 | 45.45 | AF-P31995-F1-model_v1_A |
| **KIQDSL** | Phosphorylase b kinase regulatory subunit beta | 0.13 Å | -1.76 | 46.15 | AF-Q93100-F1-model_v1_A |
| **KIQDSL** | Long-chain-fatty-acid-CoA ligase 5 | 0.40 Å | -1.47 | 15.00 | AF-Q9ULC5-F1-model_v1_A |
| **VYDPL** | Actin-binding protein IPP | 0.15 Å | -1.74 | 33.33 | AF-Q9Y573-F1-model_v1_A |
| **SAIGKI** | Ran-GTP binding protein | 0.17 Å | -1.72 | 35.29 | AF-O60518-F1-model_v1_A |
| **LPDPS** | Semaphorin 7a | 0.63 Å | -1.23 | 7.94 | AF-O75326-F1-model_v1_A |
| **VLYNS** | U2 snRNP-associated SURP motif-containing protein | 0.20 Å | -1.69 | 25.00 | AF-O15042-F1-model_v1_A |
| **KLPDD** | F-box only protein 3 | 0.28 Å | -1.60 | 17.86 | AF-Q9UK99-F1-model_v1_A |
| **NLLLQ** | Ankyrin 3 | 0.47 Å | -1.40 | 10.64 | AF-Q12955-F1-model_v1_A |
| **DNTFV** | N-acetylgalactosamine-6-sulfatase | 0.47 Å | -1.40 | 10.64 | AF-P34059-F1-model_v1_A |

**Table S3.** Number of pentapeptide pairs per pentapeptide identity / secondary structure category for both analyses (with and without parent sequence identity filter).

| Pentapeptide<br>% Identity | Number of Pentapeptide Pairs |  |  |  |  |  |
| --- | --- | --- | --- | --- | --- | --- |
|  | No Parent Identity Filter |  |  | 30% Parent Identity Filter |  |  |
|  | Helix | Extended | Coil | Helix | Extended | Coil |
| 0 | 962 | 962 | 954 | 1999 | 2000 | 1999 |
| 20 | 964 | 967 | 955 | 1999 | 2000 | 1999 |
| 40 | 967 | 966 | 954 | 1999 | 1999 | 1999 |
| 60 | 965 | 969 | 953 | 1999 | 2000 | 1999 |
| 80 | 966 | 966 | 953 | 1999 | 1992 | 1998 |
| 100 | 944 | 925 | 903 | 1508 | 921 | 1236 |

**Fig S1.**
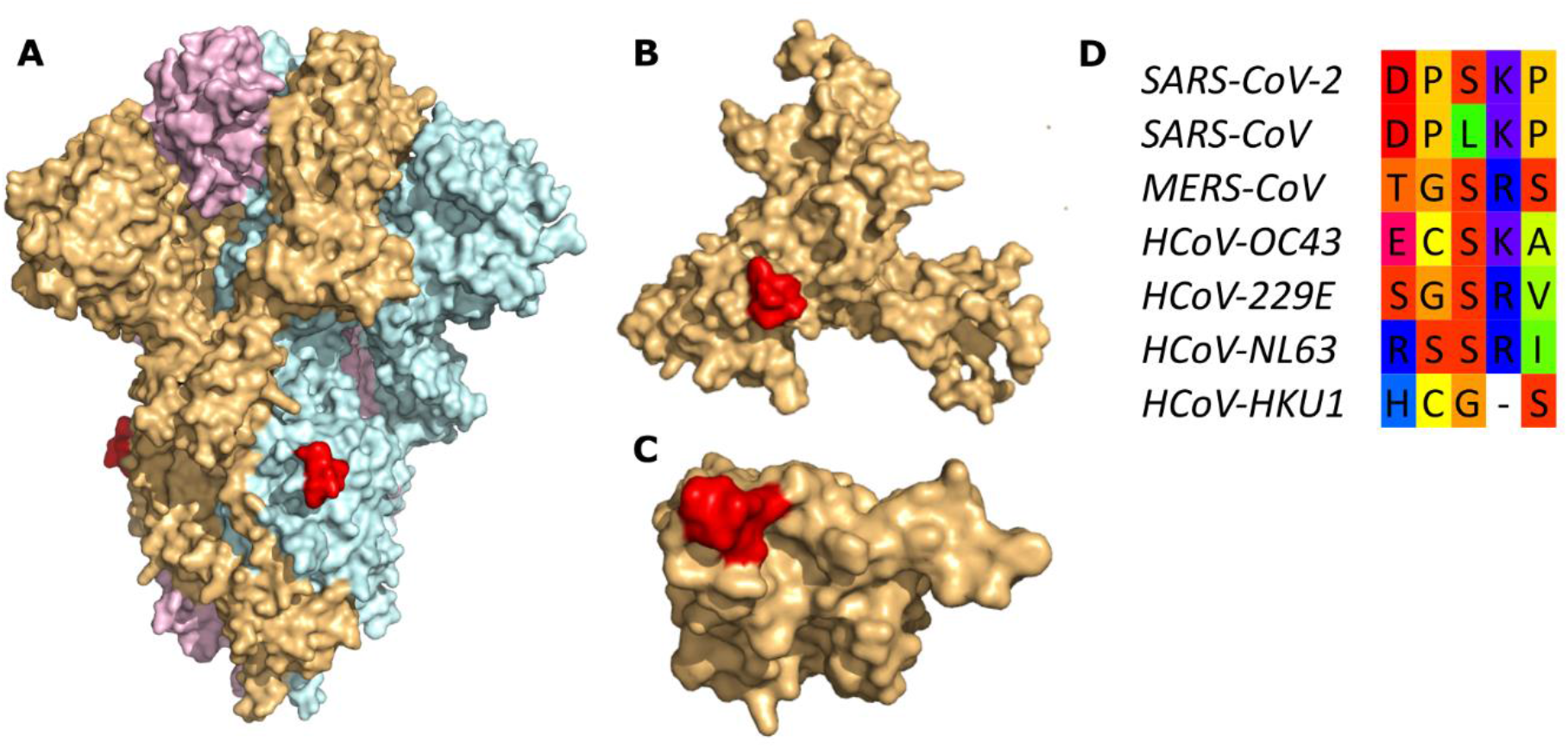
The molecular mimicry motif DPSKP (red) from Spike (A, colored by chain) matches ribosomal protein L3 (B, beige) from *Homo sapiens* with an RMSD of 0.09 Å and alanine and proline-rich secreted protein apa precursor (C, beige) from *Mycobacterium tuberculosis* with an RMSD of 0.22 Å. The motif is not conserved in human betacoronaviruses (D). Protein structures visualized can be found in Table S1. Sequences for human betacoronavirus Spike proteins were aligned using MAFFT. The molecular mimicry motif region was extracted from the alignment according to Table S1. Accessions for the sequences in order of appearance are: YP_009724390, YP_009825051, YP_009047204, YP_009555241, NP_073551, YP_003767, YP_173238.

**Fig S2.**
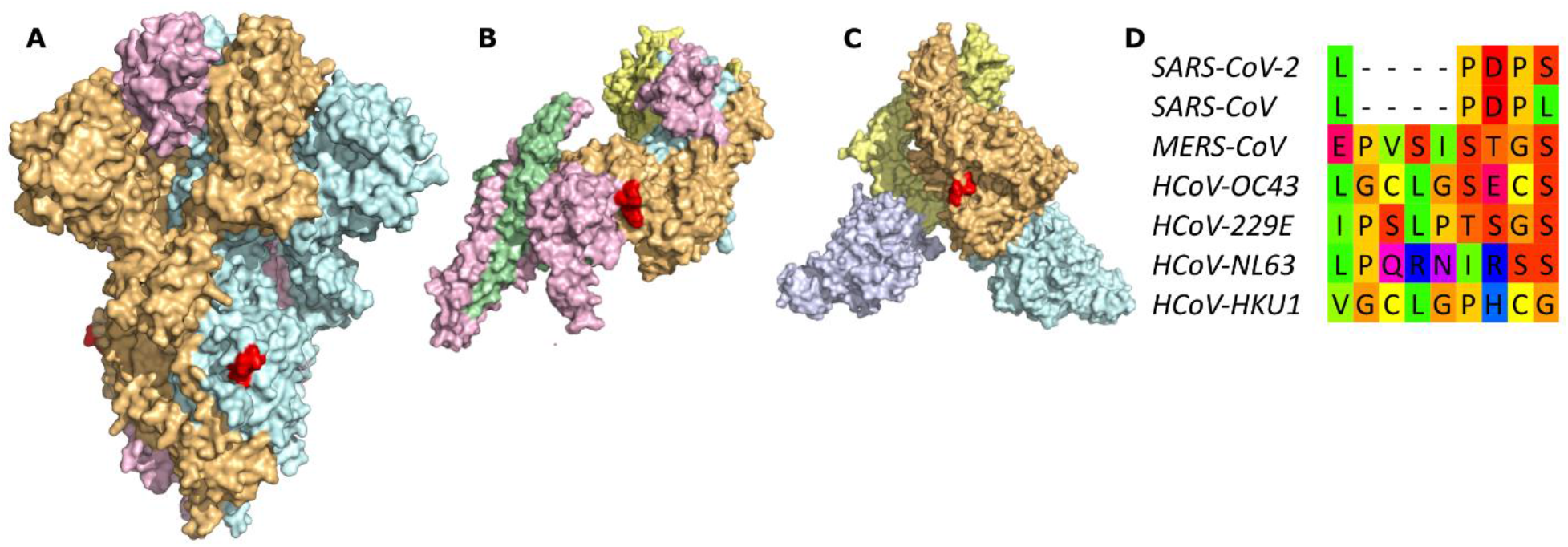
The molecular mimicry motif LPDPS (red) from Spike (A, colored by chain) matches BRCA1-A complex subunit BRE (B, colored by chain) from *Homo sapiens* with an RMSD of 0.18 Å and semaphoring-7A (C, colored by chain) from *Homo sapiens* with an RMSD of 0.66 Å. The motif is not conserved in human betacoronaviruses (D). For details, see legend of Fig S1.

**Fig S3.**
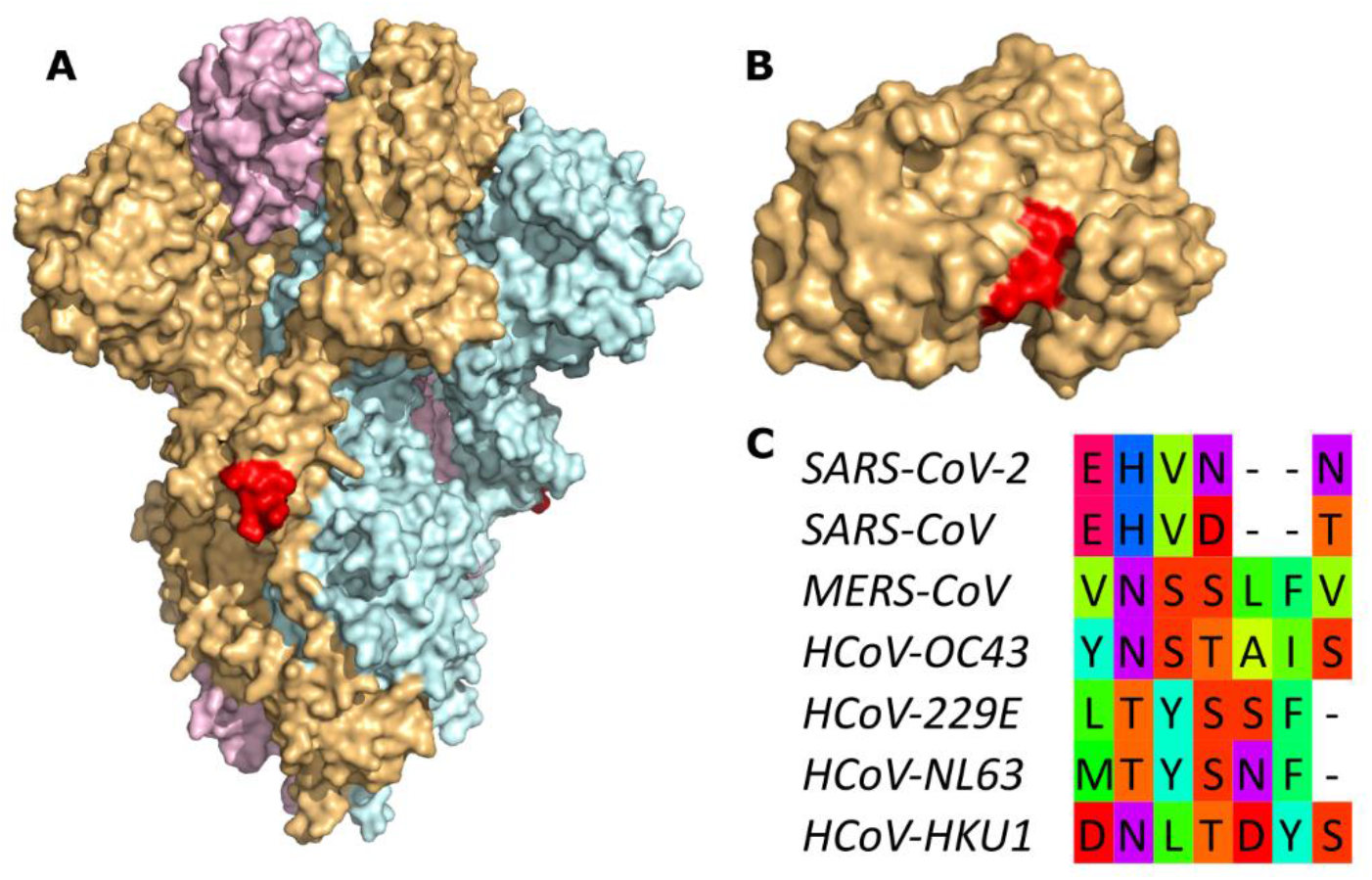
The molecular mimicry motif EHVNN (red) from Spike (A, colored by chain) matches casein kinase 2 alpha isoform (B, beige) from *Homo sapiens* with an RMSD of 0.30 Å. The motif is not conserved in human betacoronaviruses (C). For details, see legend of Fig S1.

**Fig S4.**
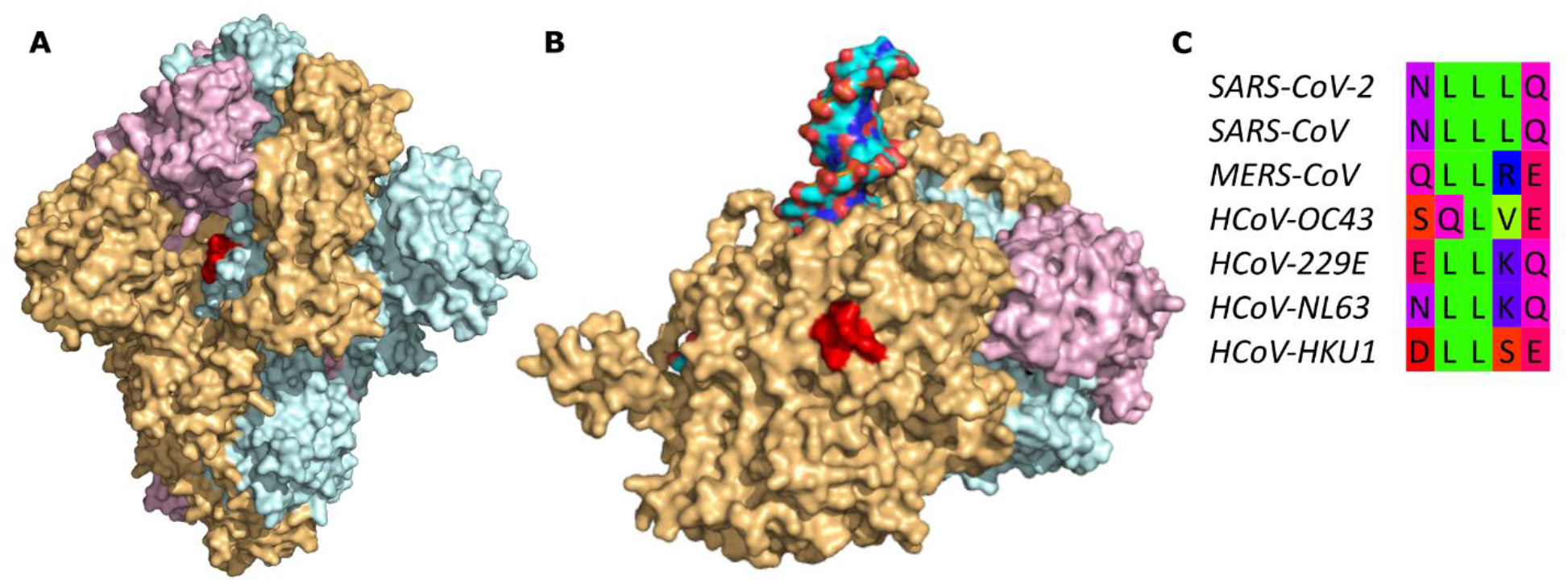
The molecular mimicry motif NLLLQ (red) from Spike (A, colored by chain) matches DNA polymerase subunit gamma-1 (B, colored by chain, with DNA colored by element) from *Homo sapiens* with an RMSD of 0.42 Å. The motif is semi-conserved in human betacoronaviruses (C). For details, see legend of Fig S1.

**Fig S5.**
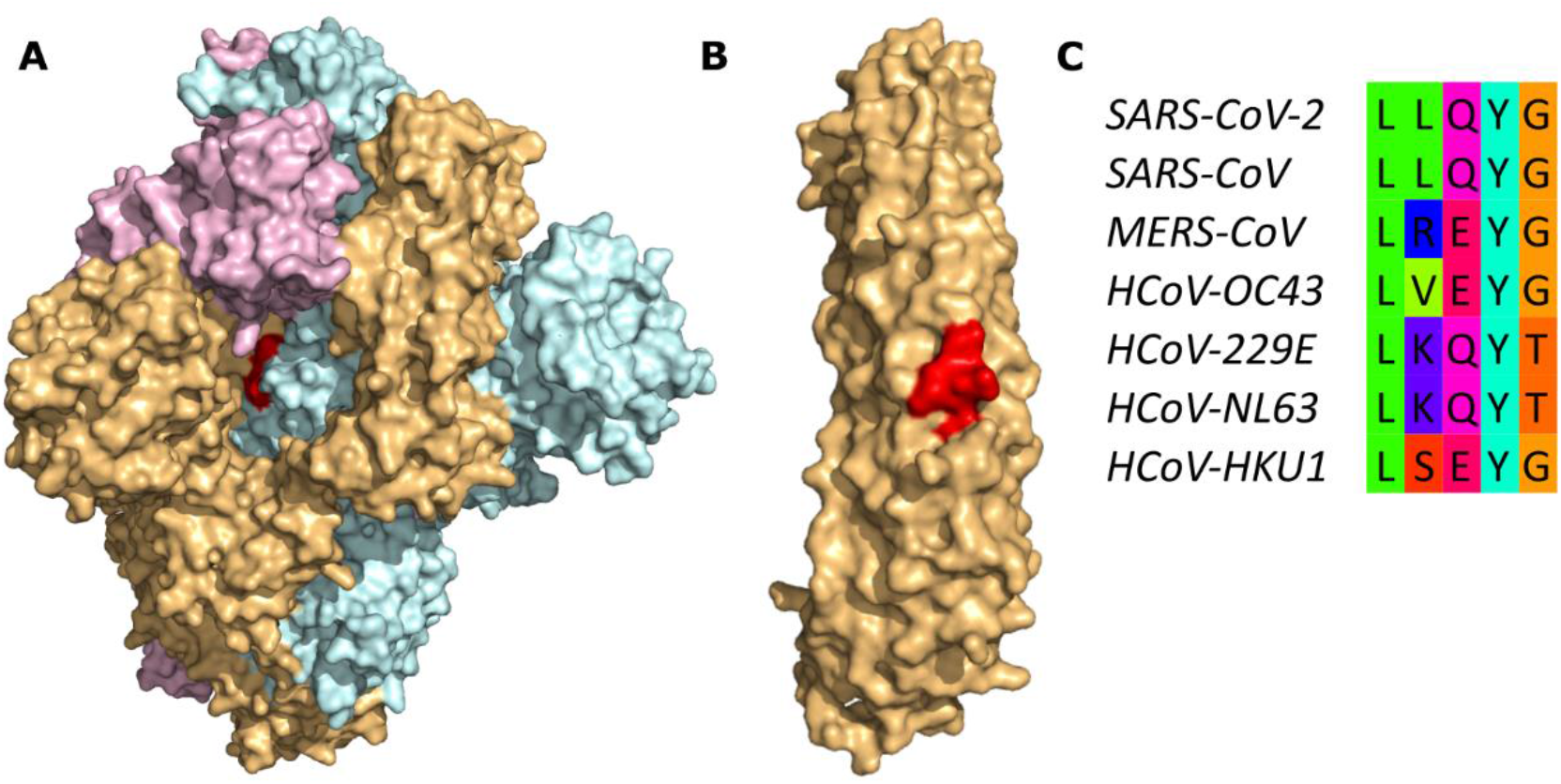
The molecular mimicry motif LLQYG (red) from Spike (A, colored by chain) matches ankyrin-1 (B, beige) from *Homo sapiens* with an RMSD of 0.49 Å. The motif is semi-conserved in human betacoronaviruses (C). For details, see legend of Fig S1.

**Fig S6.**
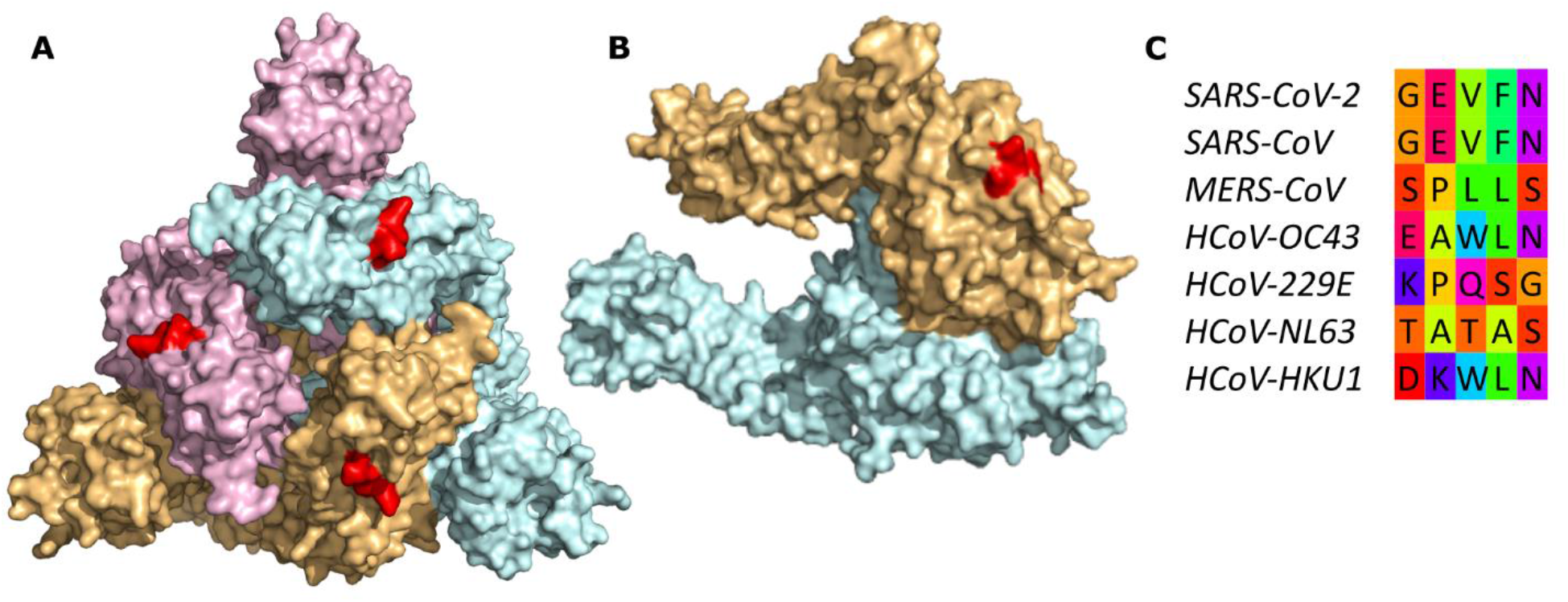
The molecular mimicry motif GEVFN (red) from Spike (A, colored by chain) matches integrin beta-1 (B, colored by chain) from *Homo sapiens* with an RMSD of 0.67 Å. The motif is not conserved in human betacoronaviruses (C). For details, see legend of Fig S1.

